# ortho_seqs: A Python tool for sequence analysis and higher order sequence–phenotype mapping

**DOI:** 10.1101/2022.09.14.506443

**Authors:** Saba Nafees, Venkata Naga Pranathi Vemuri, Miles Woollacott, Ahmet Can Solak, Phoenix Logan, Aaron McGeever, Olivia Yoo, Sean H. Rice

## Abstract

**Motivation:** An important goal in sequence analysis is to understand how parts of DNA, RNA, or protein sequences interact with each other and to predict how these interactions result in given phenotypes. Mapping phenotypes onto underlying sequence space at first- and higher order levels in order to independently quantify the impact of given nucleotides or residues along a sequence is critical to understanding sequence–phenotype relationships.

**Results:** We developed a Python software tool, ortho_seqs, that quantifies higher order sequence-phenotype interactions based on our previously published method of applying multivariate tensor-based orthogonal polynomials to biological sequences. Using this method, nucleotide or amino acid sequence information is converted to vectors, which are then used to build and compute the first- and higher order tensor-based orthogonal polynomials. We derived a more complete version of the mathematical method that includes projections that not only quantify effects of given nucleotides at a particular site, but also identify the effects of nucleotide substitutions. We show proof of concept of this method, provide a use case example as applied to synthetic antibody sequences, and demonstrate the application of ortho_seqs to other other sequence–phenotype datasets.

**Availability:** https://github.com/snafees/ortho_seqs & documentation https://ortho-seqs.readthedocs.io/

## 1 Introduction

Biological sequences (DNA, RNA, and protein) are complex molecules, consisting of many parts that interact nonlinearly with one another to give rise to output phenotype. Sequence covariation and interaction are integral parts of the structure and function of many types of RNAs and proteins (22, 21). Thus, one important task at the interface of molecular biology and bioinformatics is to pursue a holistic understanding of sequence information and how it impacts sequence function. While there exists a plethora of computational and machine learning methods to analyze sequence statics and structure, the problem of mapping phenotypes to sequence space has not been analytically explored in depth (10). We define phenotype to be any characteristic of a given sequence that can be measured, such as binding affinity of a protein to the corresponding binding site. There also continues to be rapid growth in the development of computational and machine learning approaches that attempt to not only analyze sequence structure, but also connect phenotype to sequence information (2, 1). Though many of these approaches are powerful tools, they lack the ability to quantify higher order interactions in a way that captures independent effects on the output phenotype of having given nucleotides at specific sites across a sequence. When machine learning models are used in this space, it can be difficult to extract information due to the “black box” nature of what the algorithms learned. Thus, their utilization comes with interpretability costs (28, 3).

Here, we introduce a python tool, ortho_seqs, that uses our previously described method of multivariate tensor-based orthogonal polynomials to quantify and predict first- and higher order sequence–phenotype interactions. This is done by building a sequence space given DNA, RNA, or protein sequence data and mapping corresponding phenotypes onto the space (16, 20). Projecting corresponding phenotype data onto the sequence space allows us to quantify first- and higher order relationships between sequence and output phenotype (Figure 1). For first order analysis, phenotype information is mapped onto the sequence space to quantify the effects of having a specific nucleotide or amino acid at a given site, independent of any interactions this nucleotide might have with another nucleotide at another site. For second order analysis, phenotype information is mapped onto pairs of nucleotides present on pairs of sites independently of first order effects. Third order polynomials are used to map phenotypes onto combinations of three sites and so forth. This package is written as a command-line interface that allows users to input a file with their sequence and phenotype data. It is accompanied by a graphical user interface (GUI) to provide broader access to the community.

**Fig. 1.**
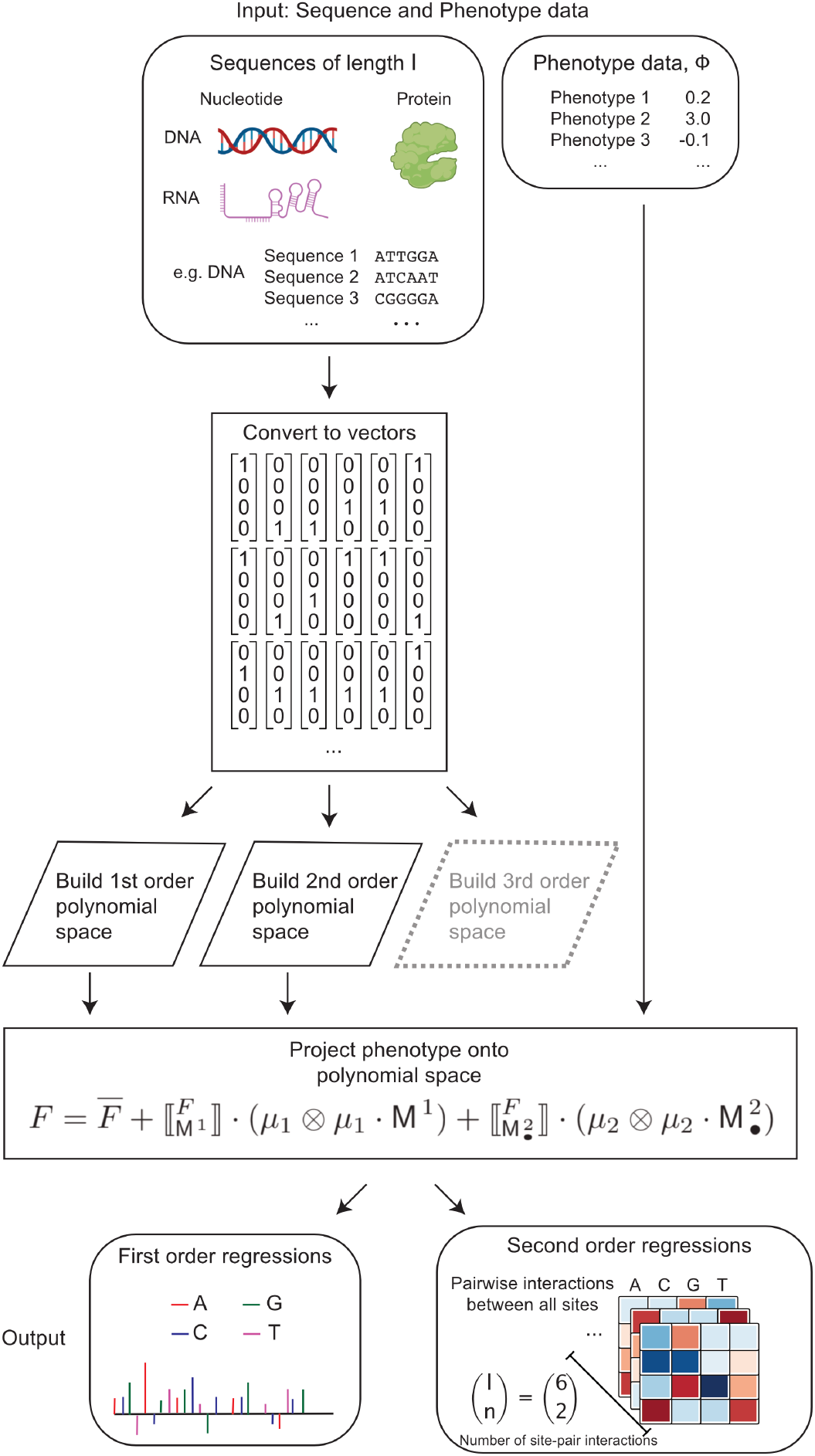
Flow chart of various parts of ortho_seqs. The tool takes user provided sequence and phenotype information as inputs, converts each nucleotide/amino acid in each sequence into n-dimensional vectors, computes various parameters such as mean, variance, and covariances, and finally builds the polynomial space based on these vectors. The second part of the tool takes the phenotype information and projects it onto the polynomial space. These projections (or regressions) are plotted as bar plots and heatmaps and saved as .png files.

## 2 Methods

The basic outline of the method can be divided into two parts (Figure 1). The first part consists of transforming the sequence data into vectors and subsequently building a multivariate tensor-based orthogonal polynomial space as shown in previous work that introduced this method (16, 20). The second part consists of projecting corresponding phenotypes, defined as real numbers and provided by the user, onto first and second order orthogonal polynomials built in the first part of the program. The projections, which are equivalent to regressions, onto the first order conditional polynomial tell us the relative functional importance of having a specific nucleotide or amino acid at a given site, *independent* of any underlying correlations this nucleotide might have with another nucleotide at another site. When projecting phenotypes onto the second order polynomial, we can identify *pairs* of nucleotides that are functionally important, independent of first order relationships. A more complete version of this method defines the presence of nucleotides at sites along a sequence as states and allows for the prediction of substitution effects (quantifying the effect that substituting a nucleotide for another one has on the phenotype). See Supplementary Methods on our derviation of a Conditional Regression Theorem for Sequence Traits for full details. The implementation of this complete method into the codebase is currently in development.

The first part of the program computes some important quantities as follows:

- **Mean**: This array tells us the mean of a specific nucleotide or amino acid at a given site across the population of sequences. This can be visualized via logo plots by using the *logo-plot* function in the command-line tool. See online documentation at https://ortho-seqs.readthedocs.io/en/master/logo_tutorial.html.
- **Variance**: This array contains the variance of a given nucleotide or amino acid at each site. The amount of variation that exists at a given site across the length of the sequence is an important characteristic of a population of sequences.
- **Covariance**: The matrix of covariance for a pair of two sites is the mean, across all individuals, of the outer product of 𝕄^1^ and 𝕄^2^, where 𝕄^1^ and 𝕄^2^ are first order vectors for each individual in the population:

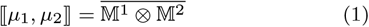 The matrices computed here are matrices of covariances (instead of the traditional covariance matrix) for a given population of sequences. This tells us which nucleotide or amino acid at a given site tends to co-occur with another nucleotide or amino acid at another site. The program outputs a .csv file of all these covariances, along with information on what nucleotides at a given pair of sites correspond to a given covariance value (Figure 3a). It also outputs a histogram of the distribution of all these values (Figure 3b). The tail ends of this distribution reveal pairs of nucleotides that covary highly with each other, either positively or negatively. These pairs are likely to be of interest given the question at hand. For example, this approach has been utilized to reveal interactions across synthetic RNA sequences (16) and can be used to infer secondary structure information (21). Figure 3 shows an example of the covariances obtained from applying the tool to analyze synthetic antibody sequence and phenotype data.
- **Regressions of phenotype onto first order polynomial**: This is the main result when running a first order analysis on a set of sequences with corresponding phenotypes (denoted in the program as *rFon1D*). These are the regressions, or projections, of the phenotype onto each site and each nucleotide or amino acid present at the site (onto the first order conditional polynomial). This can be visualized as barplots or heatmaps, where one axis represents the sites across a sequence and another represents the regressions. These plots are provided as outputs after a successful ortho_seqs run (see Figures 4–6 as examples). The distributions of these regressions can also be plotted, regardless of position along the sequence, as box plots or violin plots, as shown in the second part of Figures 4–6. Barplots, heatmaps, boxplots, and a histogram of regressions (with kernel density curve) are all saved after the *rf1d-viz* command is run.
- **Regressions of phenotype onto second order polynomial**: These are regressions of the phenotype onto a *pair* of sites, independent of first order effects (denoted in the program as *rFon12*). This is the main result when doing second order analysis on a set of sequences with corresponding phenotypes. Since these regressions are on pairs of sites and each site has a given nucleotide or amino acid present at the site, they are best represented as heatmaps (see Figure 1 and online documentation).

## 3 Implementation

ortho_seqs can be run on the command-line or through the GUI with sequences and phenotypes as inputs (Figure 2). The following sections summarize the main components and results.

**Fig. 2.**
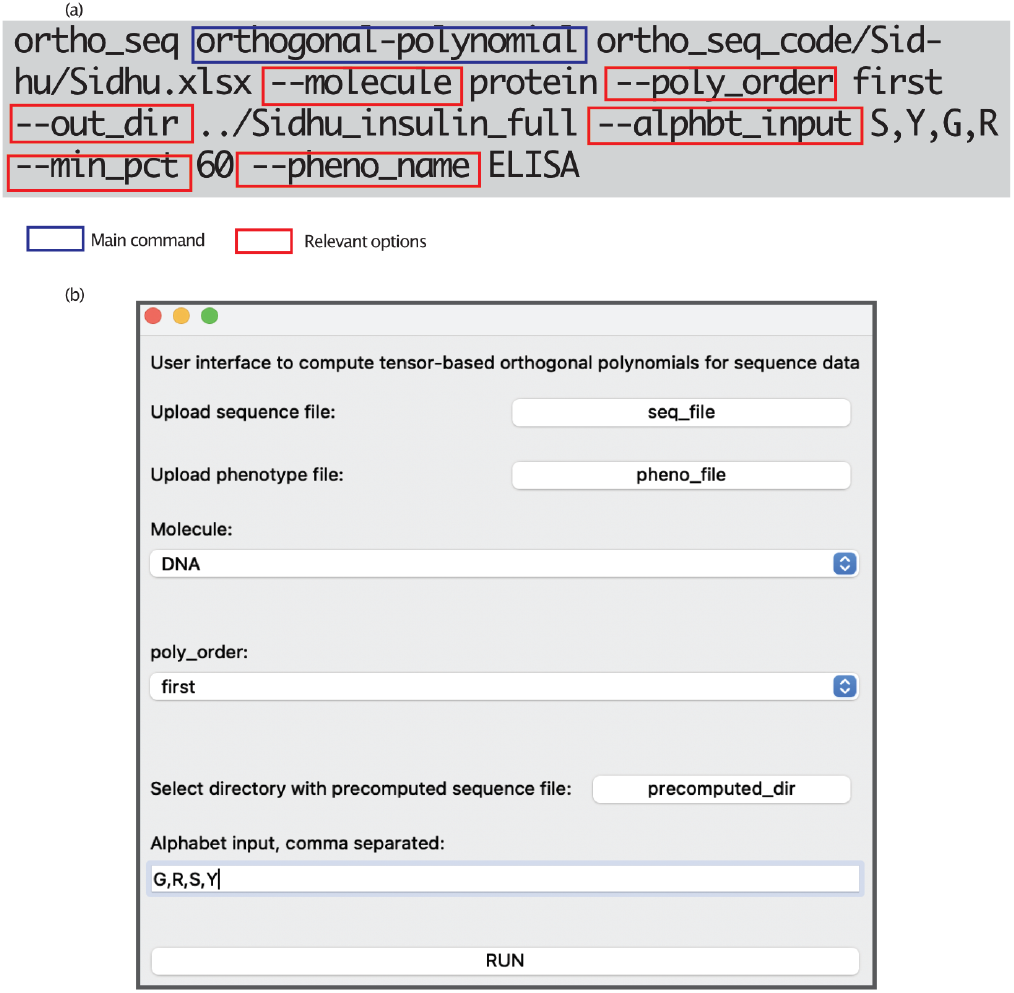
(a) ortho_seqs command-line interface showing the various flags and parameters. The main command of the tool is the *orthogonal_polynomial* command, which constructs first- and higher order tensor-valued orthogonal polynomials based on sequence data. Following this command, the user inputs a file containing sequence and phenotype information. (b) ortho_seqs GUI allows for an alternative way to run the tool and for quick exploration of sequence data..

### 3.1 Input sequence types

The tool takes in DNA, RNA, and protein sequences as inputs. The user can specify the input molecule given their datatype. Sample use cases and test datasets for each molecule categories are given in the repository and documentation.

### 3.2 Sequences of unequal lengths

The program takes in sequences of equal and unequal lengths. It automatically adds a stretch of lowercase *n*’s to sequences of unequal lengths at the end. If users wish to have padding elsewhere in the sequence, they must provide sequences with lowercase *n*’s already included. Analyzing sequences of unequal lengths is critical because in many cases, not all naturally occurring biological sequences are of equal length. For example, in the case of antibodies, the CDRH3 region (complementarity-determining region 3 of the heavy chain) contains the most diversity and is most responsible for binding to a target antigen (27). These sequences, or loops in 3D space, can vary greatly in length. Some studies have shown that the sequence length of this region matters and in some cases, shorter CDR3 loops indicate antigen exposure (8). This is indeed relevant to the current pandemic as studies have suggested that for antibodies against SARS-CoV-2, the length of CDRH3 loops plays a role in binding to the virus (23).

### 3.3 Encoding of sequence information

When the user inputs a set of sequences (nucleotide or protein), they have an option to group the various alphabets together based on biochemical properties or any other user-defined grouping or encoding (Supplementary Figure S1). For example, this is important for cases where the user wants to only probe the impact of a specific grouping of amino acids (see use case example in Results). When alphabets in amino acid or nucleotide sequences are grouped this way, the vectors for each alphabet at each site are of lower dimensions, thus also saving computational resources.

### 3.4 Protein encodings

Given protein sequences as inputs, users can utilize built-in protein encodings that are based on reduced amino acid definitions. For example, if we’re interested in breaking our amino acids into groups based on their hydrogen bonding capacity, we can use the *–alphbt_input HBOND* flag in the command-line input (Supplementary Table 1). Similarly, the amino acids in a given dataset can be broken into groups based on their hydrophobicity levels by using the *–alphbt_input HYDROPHOBICITY* flag which allows us to divide residues into four categories (15). Furthermore, when studying amino acid composition of CDRH3 loops of antibodies, certain residues, such as tyrosine (Tyr), appear more often than others depending on the length of the CDRH3 loop (9). Supplementary Table 1 contains the current list of built-in groupings within ortho_seqs.

### 3.5 Covariances of nucleotides/amino acids at different sites

ortho_seqs allows the user to analyze the statics of the sequence space by computing parameters such as mean, variance, and covariances of two states at two sites. The tool outputs a plots the distribution of the magnitudes of covariances that can help identify pairs of nucleotides or amino acids at different sites that are covarying highly positively or negatively. For sequences with a large number of sites and many states, the number of covariances can be very large. To deal with this, the user can set a value for the *–min_pct flag* to only capture the covariances in a given percentile (e.g., 90% or above). This information is also captured in a .csv file and saved in the results folder (Figure 3a).

**Fig. 3.**
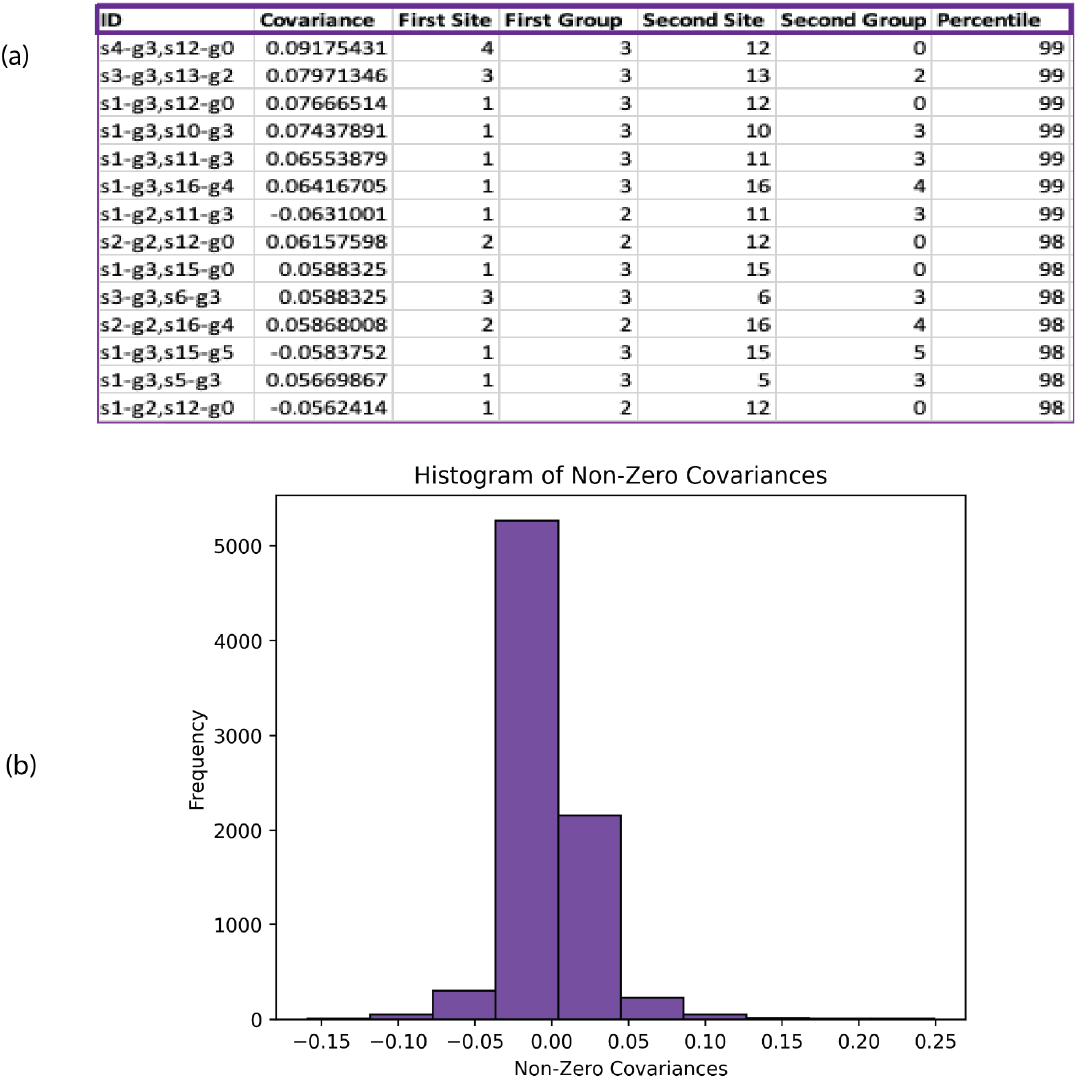
Covariance outputs from ortho_seqs for CDRH3 regions of Fabs selected for binding to insulin as the antigen. (a) A snippet of the .csv produced by ortho_seqs that contains covariance values, the pairs of sites that correspond to a given covariance value, and the amino acids present at those sites. The groups refer to user-defined encodings, which could be nucleotides or amino acids. In this case, the first four groups are the 4 amino acids present in the synthetic antibody dataset (G,R,S,Y, which are denoted as groups 0, 1, 2, and 3, respectively) in the Fab library, the fifth group is ‘z’, which denotes every other amino acid that appears in the sequence dataset, and the sixth group is ‘n’, which denotes a gap at the end of a sequence because not all CDRH3 sequences are of the same length. (b) A histogram produced by the tool, showing the distribution of non-zero covariances.

### 3.6 Sequence–phenotype mapping

Once the tool constructs the tensor-based orthogonal polynomial space based on the sequence inputs, the corresponding phenotypes are projected onto the sequence space. This projection of the phenotype onto each site is given by the *rFon1D* variable that is plotted as a bar plot and saved in the user-specified output directory. These regressions are on the first order conditional polynomial, which quantifies the effect of the phenotype onto each site and each nucleotide, amino acid, or encoding independently of any correlations that the nucleotide at that site has with any other nucleotide at another site along the sequence. These first order regressions can be compared with the outputs of the covariance analysis to see if highly covarying sites also had an impact on the structure and functionality of the sequence.

### 3.7 Higher order analysis

Currently, the tool builds up to second order tensor-based orthogonal polynomials based on sequence information and projects phenotypes onto the sequence space. The projections of the phenotype onto the first order conditional polynomial (denoted as *rFon1D*) tells us the functional importance of each site, independent of any correlations it might have with another site, as well as the functional importance of the given nucleotide or amino acid on that site. The projections of the phenotype onto the second order conditional polynomial quantifies the effect of *pairs* of nucleotides at *pairs* of sites, independent of other pairs. Furthermore, these second order effects are independent of first order relationships. Similarly, third order analysis quantifies the effect of phenotype at a combination of three sites and a given triplet of nucleotides at those sites (to be included in an upcoming release). This type of analysis can aid in the identification of sequence motifs such as those bound by transcription factors and RNA-binding proteins (19, 14).

### 3.8 Projecting different phenotypes onto the same sequence space

Given different phenotypes corresponding to the same set of sequences, the user needs to build the polynomial space based on sequence data only once. After this space has been constructed, a different phenotype can be projected onto the same space using the *–precomputed* flag. For example, in the application of ortho_seqs to the synthetic antibody dataset used in this paper, ELISA (enzyme-linked immunosorbent assay) specificity signals were measured for a set of antigen-binding Fabs that were selected to bind to three different antigens (VEGF, insulin and HER2). In this case, once the sequence space based on the CDRH3 loops was constructed, ELISA values for the cognate and non-cognate antigens were projected onto the same space to quantify the effect of specific and non-specific binding, as shown in the Results section.

### 3.9 Visualizing results

Once a successful first order run is completed, all output result files are stored in the user-defined output directory. Some of the plots that are generated after a successful run include a histogram depicting covariances of nucleotides at different sites (as described in the covariances section above) and first order regressions of the phenotype onto sequence space. Furthermore, the user can run the *rf1d-viz* command to visualize these regressions in different ways, such as plotting a barplot, a heatmap, a boxplot, and a distribution of the regressions (with kernel density curve). The user can also obtain a summary of the top regressions (see detailed tutorial for examples). All resulting plots are saved as high quality images (.png files), which can then be used for publication or reports (see examples shown in Figures 4-6 and Supplementary Figures S2-S4).

**Fig. 4.**
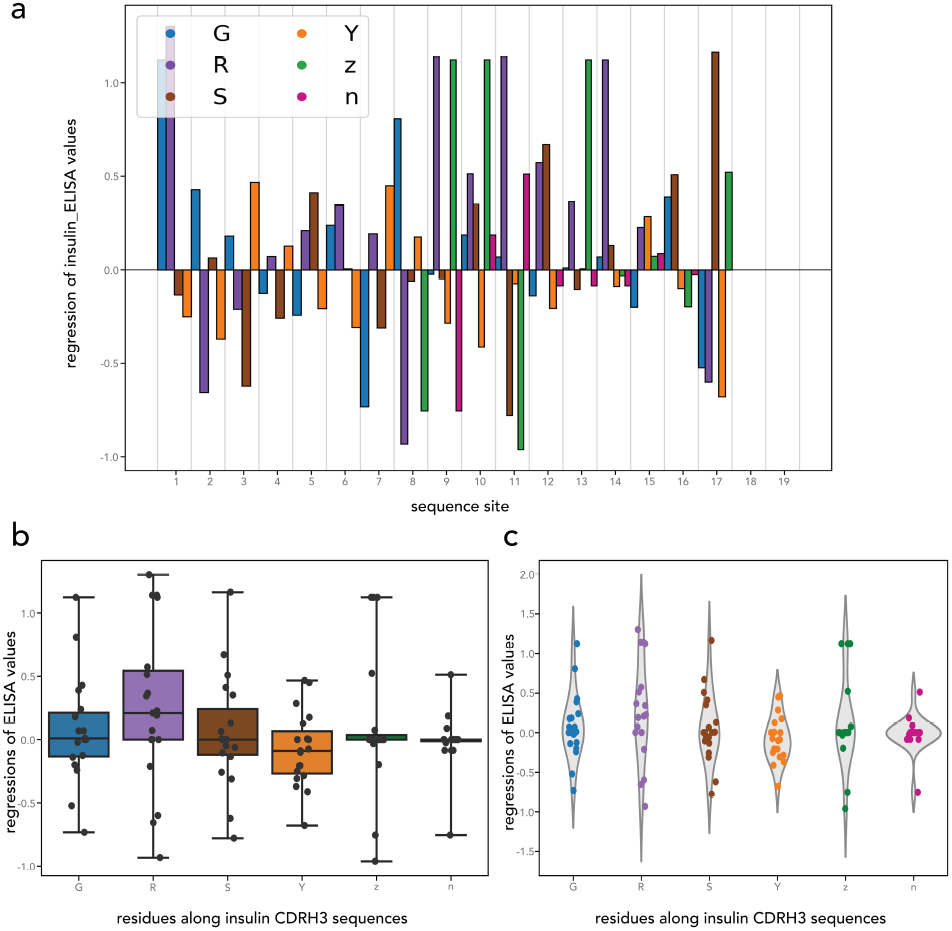
Projecting insulin ELISA specificity values onto CDRH3 sequences of Fabs designed to bind to insulin to assess the effects of residues on antigen specificity. Here, z refers to other residues that appeared in the sequence dataset (A,F,I,L,M). (a) Barplot showing regressions of ELISA specificity values onto each site and each of the given residues across the CDRH3 loops of the synthetic Fabs binding to insulin (target antigen). This can also be showcased as a heatmap, along with a distribution of the regressions as a histogram (Supplementary Figure S2). (b) Box plots and (c) violin plots showing distributions of regressions for each residue regardless of location in the CDRH3 loop.

## 4 Results

### 4.1 Application to synthetic antibody sequences

To show proof of concept of this method as applied to protein sequences, we used previously published synthetic antibody sequence data and corresponding phenotypes from 4, henceforth referred to as the *synthetic antibody dataset*. In this work, the authors constructed synthetic antibody libraries to measure the impact of four different amino acids, tyrosine (Y), serine (S), glycine (G), and arginine (R) on antigen recognition. They tested three different antigens, insulin, VEGF, and HER2. Affinity and specificity data for these antigen-binding antibody fragments (Fabs) was provided for these three antigens. For sequence data, we used the CDRH3 loop sequences. For corresponding phenotype data, we used the given relative binding values given by the single-point competitive phage ELISA. The use case example shown here uses CDRH3 regions of Fab sequences selected for binding to insulin as the target antigen.

When inputting this data to ortho_seqs, we used the custom amino acid encoding feature and provided G, R, S, Y as our grouping and a cutoff of 60 as the value for “min_pct”, which outputs covariance values in the 60th percentile and stores them in a .csv file. The grouping creates 6-dimensional vectors: one dimension for each of these four amino acids, one dimension for ‘n’ that indicates a gap if there was no amino acid present (for unequal CDRH3 sequence lengths), and one dimension for any other amino acids present in the dataset (grouped as one group ‘z’). The first part of the analysis deals with looking at the statics of the sequence space. The covariance analysis reveals a small set of amino acid pairs covarying highly with one another (Figure 3). For example, Figure 3a shows that a tyrosine at site 4 covaries highly with a glycine at site 12 (with a covariance of .09). These covarying pairs of amino acids could be spatially or functionally important for binding to the target antigen, but this aspect is only assessed when the corresponding phenotypes are projected onto the sequence space.

#### 4.1.1 Regressions of ELISA specificity values onto the sequence space

The regressions (or projections) of the phenotype, ELISA binding specificity signal, onto the first order conditional polynomial quantifies the importance of a given residue at a site regardless of its correlations with other amino acids on other sites. The authors of the synthetic antibody dataset determined specificity of the Fabs “against a panel of antigens” by varying the frequencies of four amino acids (Ser, Gly, Tyr, Arg) in the CDRH3 loop and quantified the effect on binding specificity to the cognate antigen. Their experiments showed that an increase in tyrosine residues led to a decrease in nonspecific binding, while an increase in arginine residues led to increased nonspecific binding. Here, when analyzing insulin Fab sequences, we projected ELISA specificity values onto the CDRH3 loops binding to insulin as the cognate antigen. This analysis is done across the population of sequences, and the means of an amino acid at a given site are subtracted in the beginning when the sequence space is being calculated. This allows for the quantification of the impact of each residue at a given position along the CDRH3 loops regardless of the initial quantities of that residue.

Looking at the effect of a given residue at a position along the CDRH3 loop, there exist high and positive regressions of ELISA specificity values (when Fabs are bound to insulin as the antigen) onto arginine (R) starting at position 9 along the loop and until position 14 (Figure 4a, Supplementary Figure S2). This has potential impacts on the spatial organization of a given residue in the 3-dimensional space of the CDRH3 loop when binding to a cognate antigen (see Figure 2 in 4). The authors suggested that “though Arg residues contribute favorable binding energy in many protein-protein interactions,” they do so “only when located at precise locations within the interface.” For Tyr (Y), the regressions seem to be uniformly spread across the loop, in either the positive or negative direction. The plot of regressions in Figure 4a shows that Tyr, similar to the case of Arg, contributes favorably to specificity when presented at certain locations along the loop and otherwise contributes negatively to antigen specificity. When looking at the contributions of each residue as a whole (Figure 4b,c), we wanted to see if the distribution of regressions for Arg was significantly different from that of Tyr regressions. Conducting a Mann-Whitney U test, we see that for the case of specific binding, these two distributions are not very significantly different (*p*-value: 0.033) when taken as a whole, contrary to what the authors found. This underscores the importance of assessing where these residues are located along the loop, as shown in Figure 4a, and not just their overall impact on specificity for the cognate antigen. Thus, doing first order analyses with ortho_seqs not only reveals population-wide patterns of residues in a set of sequences, but also quantify location-specific impacts of amino acids on the corresponding phenotype.

#### 4.1.2 Quantifying nonspecific binding

To determine the effect of particular residues along the CDRH3 loops on *nonspecific* binding, we projected ELISA specificity signals of two non-cognate antigens (VEGF and HER2) onto CDRH3 loops of insulin Fabs. The authors of the dataset showed that when Arg is added to the loops, there exists more “nonspecific binding”, whereas the addition of Tyr to the loops leads to “lower nonspecific binding”. To quantify the effect of these amino acids on nonspecific binding, we projected ELISA specificity signals of the VEGF antigen onto insulin CDRH3 loops (Figure 5, Supplementary Figure S3) and HER2 ELISA signals onto insulin CDRH3 loops (Figure 6, Supplementary Figure S4). Figures 5a and 6a show that there exists large and positive regressions of specificity onto arginines across the loop, indicating that arginine does indeed contribute to nonspecific binding because the cognate antigen for these loops is insulin and not VEGF or HER2. In sharp contrast to the case of Arg and in accordance with the authors’ conclusions, the regressions of ELISA signals onto Tyr across the loop are also largely negative, indicating that Tyr is not preferred for nonspecific binding. For the case of VEGF specificities projected onto insulin loops, when looking at the distributions of the residues as a whole and assessing whether the distribution of Arg regressions are significantly different from the distribution of Tyr regressions, we note that indeed these two distributions are significantly different (Figure 5b,c; *p*-value for Mann-Whitney U test: 5*x*10*e −* 6). Similarly, for HER2 specificities projected onto insulin loops, the distributions of Arg regressions and those of Tyr regressions are significantly different (Figure 6b,c; Mann-Whitney U test *p*-value: 3.9*x*10*e−*6). This suggests that indeed, Arg and Tyr residues are playing starkly different roles in nonspecific binding, with Arg giving rise to more nonspecificity and Tyr contributing negatively to nonspecificity.

**Fig. 5.**
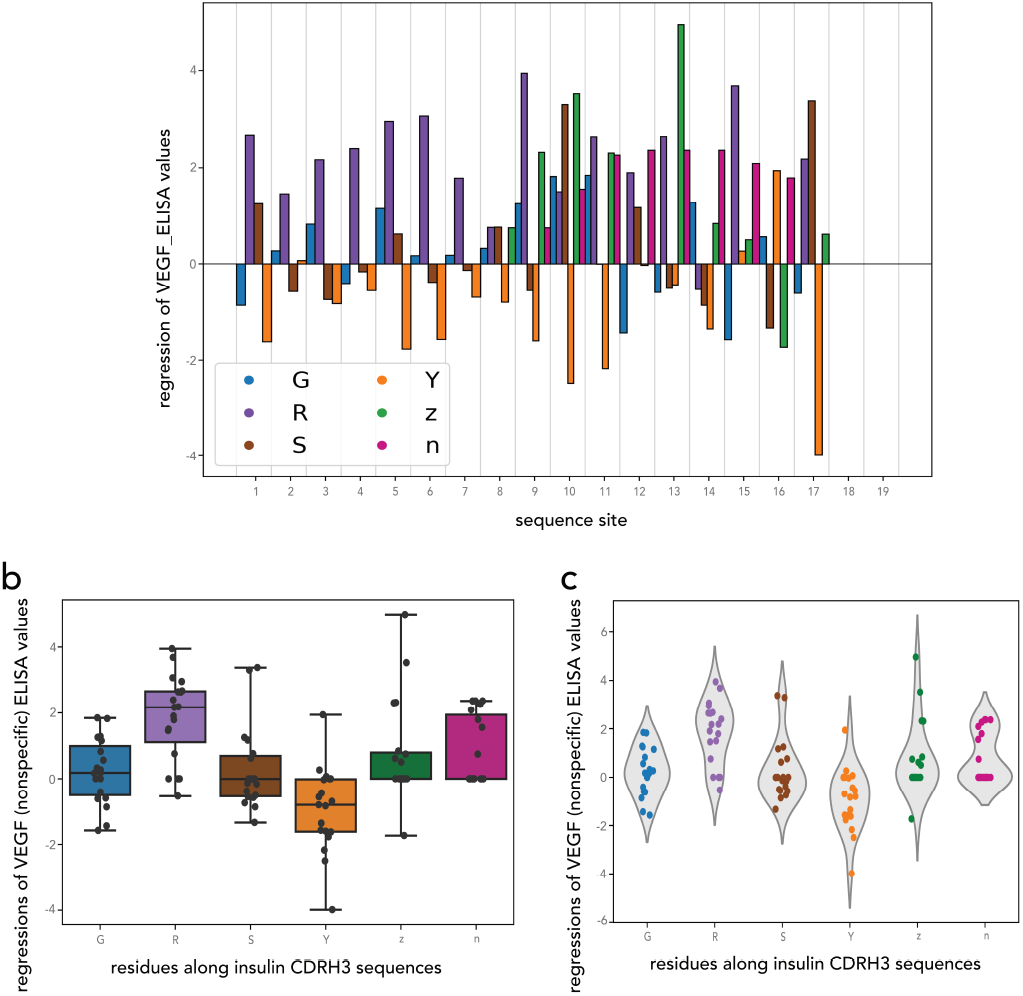
Nonspecific binding of VEGF to insulin CDRH3 sequences. Here, z refers to other residues that appeared in the sequence dataset (A,F,I,L,M). (a) Barplot showing regressions of VEGF ELISA values onto each site and each of the given residues across the CDRH3 loops of the synthetic Fabs binding to VEGF (non-target antigen). This can also be showcased as a heatmap, along with a distribution of the regressions as a histogram (Supplementary Figure S3). (b) Box plots and (c) violin plots showing distributions of regressions for each residue regardless of location in the CDRH3 loop.

**Fig. 6.**
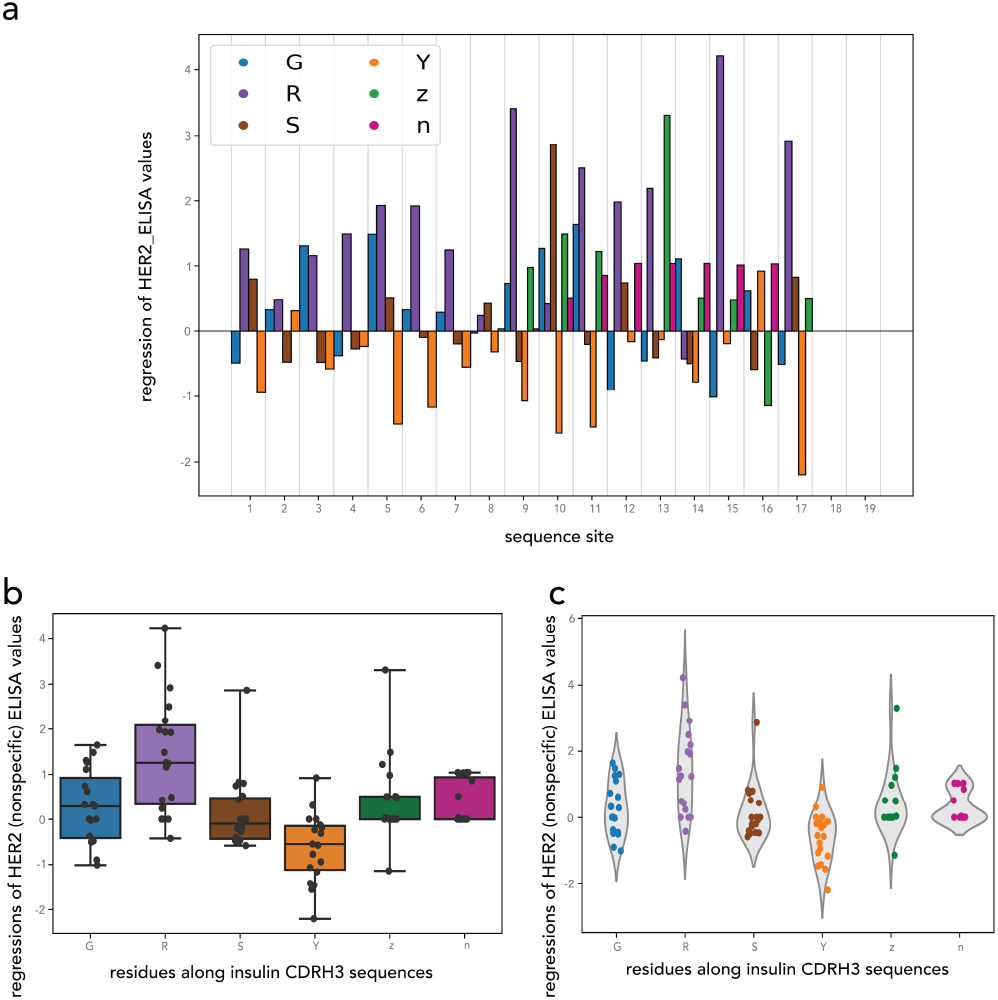
Nonspecific binding of HER2 to insulin CDRH3 sequences. Here, z refers to other residues that appeared in the sequence dataset (A,F,I,L,M). (a) Barplot showing regressions of HER2 ELISA values onto each site and each of the given residues across the CDRH3 loops of the synthetic Fabs binding to HER2 (non-target antigen). This can also be showcased as a heatmap, along with a distribution of the regressions as a histogram (Supplementary Figure S4). (b) Box plots and(c) violin plots showing distributions of regressions for each residue regardless of location in the CDRH3 loop.

To further probe the role of Arg in specific vs. non-specific binding, we compared the Arg regressions in TGMAIthe case of specific binding of insulin to insulin loops (Figure 4b,c) to the Arg regressions in non-specific binding of VEGF to insulin loops (Figure 5b,c) and to Arg regressions in non-specific binding of HER2 to insulin loops (Figure 6b,c). The distributions are significantly different between the specific and non-specific cases (*p*-values for Mann-Whitney U test: insulin regressions onto insulin loops compared with VEGF regressions onto insulin loops = 4.7*x*10*e −* 4; insulin regressions onto insulin loops compared with HER2 regressions onto insulin loops = 3.3*x*10^*−*^3).

Thus, projecting specificity signals of non-cognate antigens onto given CDRH3 loops can reveal important characteristics and preferences of residues across different parts of the loops and allows for the quantification of their impact on non-specific binding.

## 5 Discussion & Conclusions

Mapping output phenotypes onto DNA, RNA, or protein sequence space is critical in order to understand the relationship between parts of a given sequence and what functional roles these interactions play in the resulting output phenotype (12). While tremendous computational advancements have been made to understand sequence–phenotype relationships in molecular and synthetic biology, most of the current methods are machine learning and deep learning methods (17, 14, 25). Though these computational tools yield insightful results, they come at the cost of interpretability and can have considerable design constraints. Here, we have taken our previously described method of multivariate tensor-valued orthogonal polynomials for sequences and packaged it into a command-line tool, along with a GUI for wider accessibility, which can be easily utilized by users wanting to quantify the sequence–phenotype relationships in their datasets (16). To show proof of concept of the method and tool as applied to protein sequences, we used a previously published dataset of synthetic antibody sequences and quantified the positive or negative effects of certain amino acid residues along the CDRH3 loops of synthetic antibodies and how much a given residue gives rise to nonspecific and specific binding to the cognate antigen (4). In addition, the method quantifies these residue preferences at all locations across the loop, which allows us to understand the potential links between sequence composition and 3D structure, underscoring the importance of biophysical properties of various amino acids. In the case of Arginine mutations, we showed that they are preferred in a certain region (positions 9-14) of the CDRH3 loop of the insulin Fab. This is in concordance with previous studies that suggest that Arginine is a particularly “interactive” molecule, which could lead to more nonspecific interactions with the target antigen, but that these interactions are context-specific and occur in hydrophobic areas (24). Thus, information regarding location-specific effects of Arginine allows us to further probe the effect of these mutations on antibody binding given the biochemical environment occupied by these residues along the CDRH3 loop.

The ortho_seqs tool allows the user to make use of the various available built-in encodings or provide custom ones ones to focus on context-dependent biochemical properties of amino acids. For example, it has been shown that for clinical antibodies, certain amino acids are more important than others for specificity based on chemical and physical properties such as charge and whether they are acidic or basic (18). Thus, quantification of exactly how important these residues are and their location-specific interactions across the CDRH3 loop is critical to understanding antibody specificity and has implications in novel antibody design therapeutics.

Furthermore, it is increasingly apparent that sequence structure plays a large role in sequence function, underscoring the need to build methods that aim to understand sequence–structure–function relationships (7, 5). To probe the effects of structure on sequence function, ortho_seqs can be utilized to provide insights into structural patterns that arise when projecting phenotypic information onto sequence space as shown in the aforementioned case of synthetic antibody CDRH3 loops, and when identifying covariation in the case of RNA secondary structure (16, 21). In the protein space, the tool can be utilized in conjunction with existing methods to identify interacting and coevolving residues based on the underlying sequence information (11).

To leverage the advancements made in the construction of sophisticated machine learning models that aim to decipher sequence–function interactions, ortho_seqs can be used in conjunction with these prediction tools to quantify higher order interactions and generate synthetic data for further analysis. For example, complex deep learning algorithms, such as AlphaFold, have been constructed to understand interactions across protein sequences and how this affects the final output 3D structure (13, 5). Using sequences with preferred phenotypes predicted by ortho_seqs as inputs for protein modeling tools can yield powerful estimations of viable 3D structures of synthetically designed proteins. Other tools such as immuneML are used by the AIRR (Adaptive Immune Receptor Repertoires) community to generate random antibody repertoire datasets to train machine learning models, which can then be used as inputs to ortho_seqs (6, 17). The outputs from both the ML methods and the analytical approach of ortho_seqs can further inform the construction of increasingly complex classifiers to decode immune signals in antibody-antigen datasets.

## 6 Availability & Future Directions

The command-line tool and accompanying GUI is written in Python and available as a pip-installable PyPI package. This can be found at https://pypi.org/project/ortho-seq-code/1.0.1/ and on GitHub at

https://github.com/snafees/ortho_seqs. Documentation for the tool, along with extensive use case examples and tutorials are provided in readthedocs https://ortho-seqs.readthedocs.io/. Example code to run the tool is shown along with explanations of the outputs. The tool has been unit-tested and utilizes continuous integration in order to ensure that the software is working on different systems. Currently, the analysis can be done at the first and second order levels, while the third order feature will be part of the next release. Some further developmental updates will include more visualization outputs, and faster processing and parallelization to speed up runtimes. The tool can be used on high performance compute clusters, where the job can be monitored continuously via a job scheduler such as SLURM (26). The codebase is actively in development and we welcome the comments and contributions from current users and the community to facilitate further analyses and hypothesis generation.

## Acknowledgements

We thank Joan Wong, John Pak and Sandra Schmid for critical feedback and helpful discussions, Olga Botvinnik for helping with Figure 1, and Colin Zamecnik for thoughtful discussions on protein interactions.

