## Supplementary Information for "ortho_seqs: A Python tool for sequence analysis and higher order sequence–phenotype mapping"

Table 1: ortho_seqs built-in amino acid encodings for protein sequences.

| **Name of Encoding** | **Properties** | **Amino Acids** | **CLI Flag** (–alphbt_input [encoding]) |
|---|---|---|---|
| **Conventional** | - | - | CONVENTIONAL |
| | Basic | K,R,H | - |
| | Acidic | D,E | - |
| | Nonpolar | A,V,L,I,P,F,M,W,G,C | - |
| | Uncharged Polar | N,Q,S,T,Y | - |
| **Sigma's classification** | - | - | SIGMA |
| | Hydrophobic aliphatic | A,I,L,M,V | - |
| | Hydrophobic aromatic | F,Y,W | - |
| | Polar neutral | N,Q,C,S,T | - |
| | Basic | K,R,H | - |
| | Acidic | D,E | - |
| | Unique | G | - |
| | Ungrouped | P | - |
| **Hydrophobicity** (hydrophob1995) | - | - | HYDROPHOBICITY |
| | Very hydrophobic | L,I,F,W,V,M | - |
| | Hydrophobic | C,Y,A | - |
| | Neutral | T,E,G,S,Q,D | - |
| | Hydrophilic | R,K,N,H,P | - |
| **According to hydrogen bonding capability** | *Can make Hbonds* | - | HBOND |
| | Non-aromatic neutral | N,Q,S,T | - |
| | Non-aromatic charged | D,E,R,K | - |
| | Aromatic | Y,H,W | - |
| | *Can't make Hbonds* | - | - |
| | Non-aromatic | A,I,L,M,V,C,G | - |
| | Aromatic | F,P | - |
| **Frequency based amino acids in CDRH3 loops** freqaa2017 | - | - | FREQUENCY_11AA |
| | 9 most abundant | Y,G,D,V,S,A,F,R,L (each is a separate group) | - |
| | Next 6 most abundant | P,T,W,N,E,M | - |
| | Last 5 | K,I,H,Q,C | - |

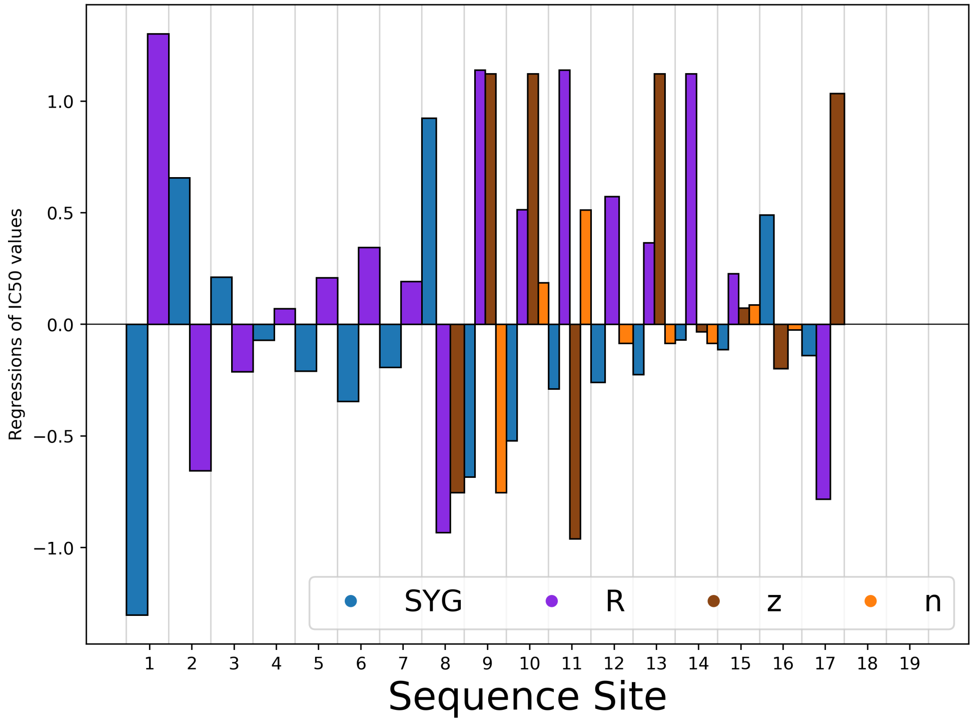


Figure 1: Regressions of insulin ELISA values onto insulin CDRH3 loops with a custom grouping of residues as inputs to ortho_seqs. Here, the residues serine (S), tyrosine (Y), and glycine (G), are grouped together while Arginine (R ) is grouped by itself. 'z' refers to other residues that appeared in the sequence dataset (A,F,I,L,M) and 'n' denotes padding at the ends of the sequences that are of unequal length. Given this grouping, the tool creates 4-dimensional vectors.

Supplementary Figures 2-4 are more plots generated by ortho_seqs depicting regressions of ELISA values of a given antigen onto insulin CDRH3 loops.

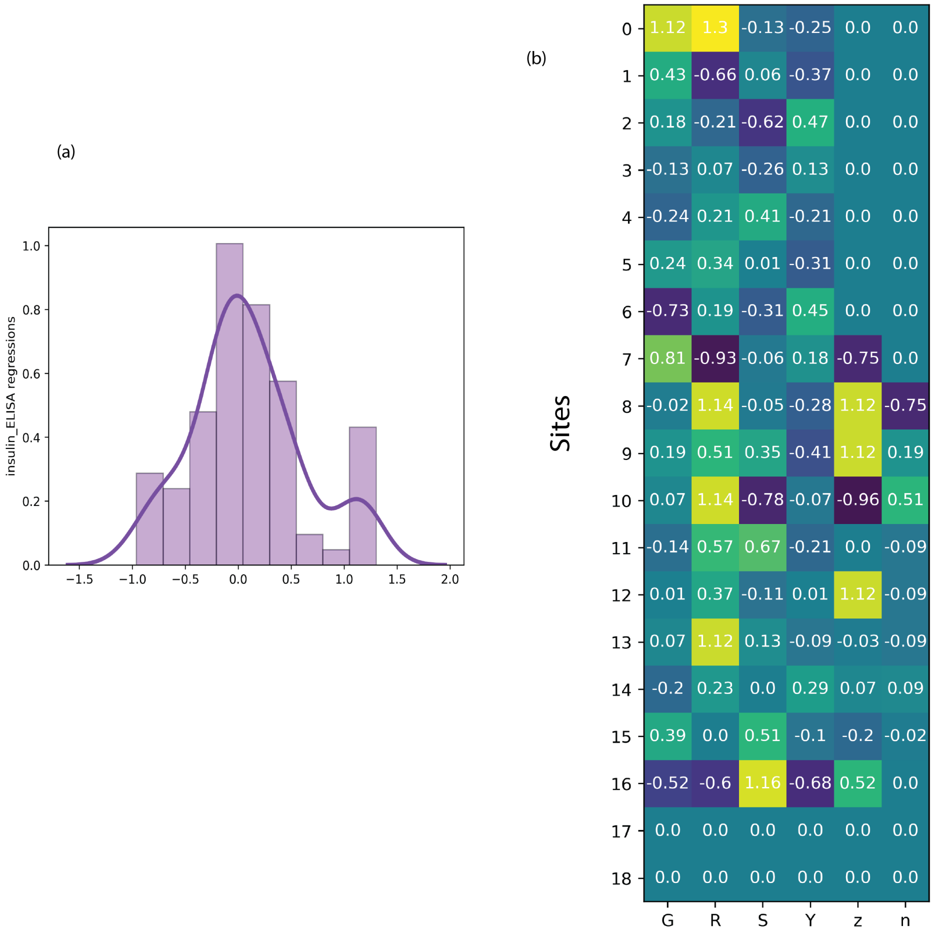


Figure 2: Regressions of insulin ELISA values onto insulin CDRH3 loops. (a) Histogram showing the distribution of the regression values along with a kernel density curve. (b) Heatmap of the regressions at each site and for each residue (same as the bar plot shown in Figure 4a).

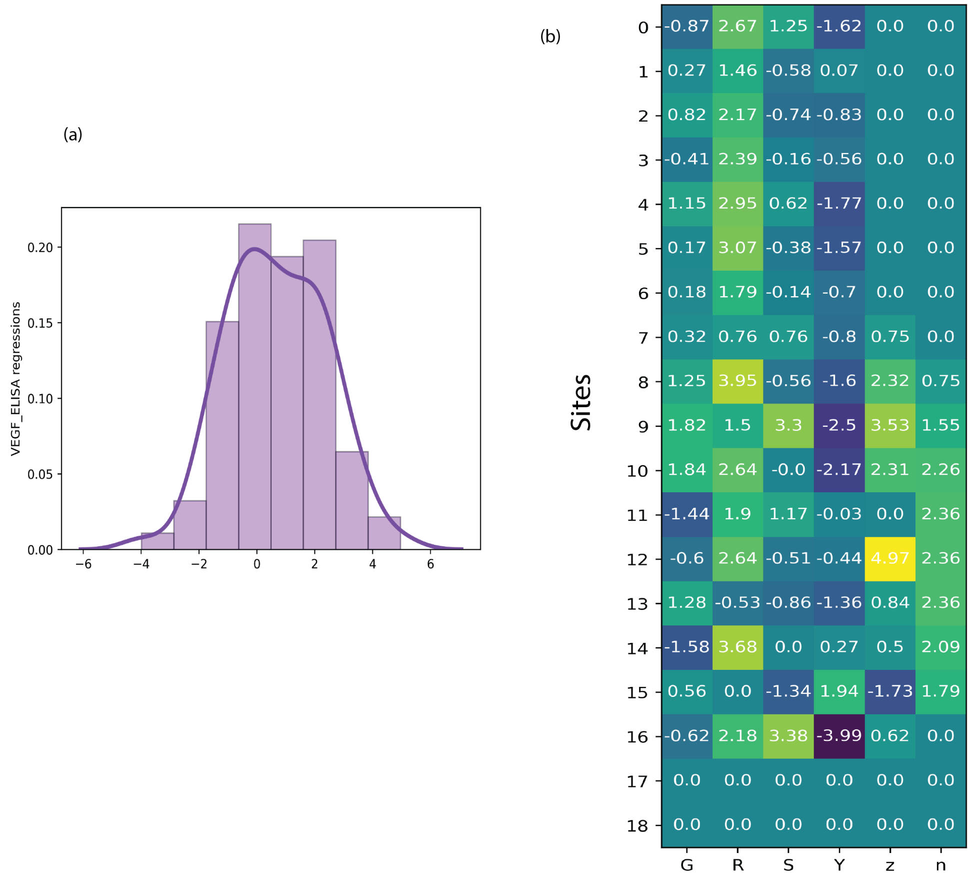


Figure 3: Regressions of VEGF ELISA values onto insulin CDRH3 loops. (a) Histogram showing the distribution of the regression values along with a kernel density curve. (b) Heatmap of the regressions at each site and for each residue (same as the bar plot shown in Figure 5a).

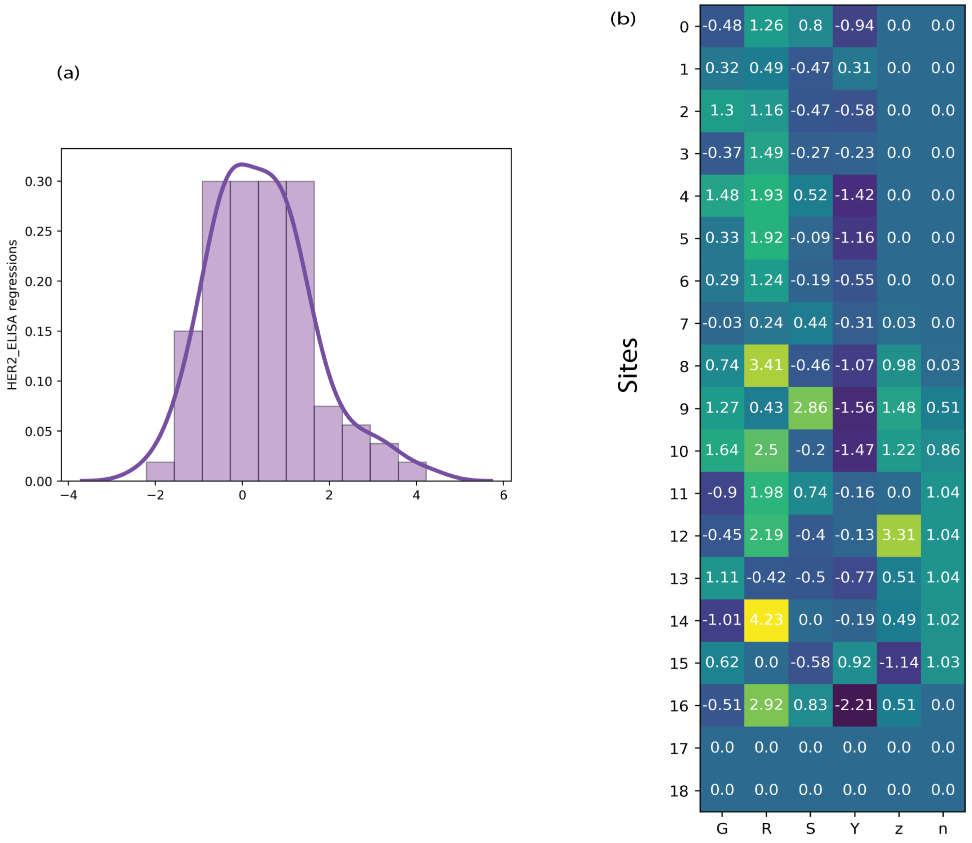


Figure 4: Regressions of HER2 ELISA values onto insulin CDRH3 loops. (a) Histogram showing the distribution of the regression values along with a kernel density curve. (b) Heatmap of the regressions at each site and for each residue (same as the bar plot shown in Figure 6a).
