## Supplementary Methods for "ortho_seqs: A Python tool for sequence analysis and higher order sequence–phenotype mapping"

### A conditional regression theorem for sequence traits

The conditional regression theorem was derived for a set of scalar traits, meaning that the value of each trait could be represented as a number. The elements of a sequence – nucleotides in DNA and RNA, amino acids in protein sequences – can not be represented as scalars because they do not differ in any obvious quantity. Instead, each site in a sequence has one of a finite number of different states, with no natural ordering of these states.

I will illustrate the approach to dealing with sequence traits using the four DNA nucleotides. The same reasoning applies to any fixed number of possible states.

Because there are only four possible, and mutually exclusive, states at a site in a DNA sequence, specifying any three of them determines the fourth (*e.g.* once we specify whether a site has an A, a C, or a T, we know whether or not it has a G). This means that we can not calculate a conditional basis as we did in section (–); if specifying C, G, and T, completely determines A, then A independent of C, G, and T is zero. This is the same problem encountered in trying to perform multiple regression on a variable with a multinomial distribution (May & Johnson, 1998).

One way of dealing with this problem in statistics is to simply ignore one of the states, since the remaining ones completely determine the state of the system. While this works mathematically, it is not acceptable in a theoretical biology context. If we were to ignore, say, T, then the effects of having a T would be spread out over the effects of the remaining nucleotides. We would not then be able to directly assess the fitness effects of having a T at a particular site, or evaluate the effects of changing a C to a T.

Fortunately, there is still a way to construct simple and conditional bases that capture each possible nucleotide, though in this case these different bases will not be biorthogonal.

#### The Simple Basis

We represent each DNA nucleotide as a vector:

$$\begin{array}{cccc} \text{A} & \text{C} & \text{G} & \text{T} \\ \begin{bmatrix} 1 \\ 0 \\ 0 \\ 0 \end{bmatrix} & \begin{bmatrix} 0 \\ 1 \\ 0 \\ 0 \end{bmatrix} & \begin{bmatrix} 0 \\ 0 \\ 1 \\ 0 \end{bmatrix} & \begin{bmatrix} 0 \\ 0 \\ 0 \\ 1 \end{bmatrix} \end{array} \tag{1}$$

Our simple basis is just the set of coordinates defined by vectors in 1 with the population means (the frequencies of each nucleotide) subtracted out.

$$
\begin{aligned}
\mathbf{P}^a &= \mathrm{A} - \overline{\mathrm{A}} \\
\mathbf{P}^c &= \mathrm{C} - \overline{\mathrm{C}} \\
\mathbf{P}^g &= \mathrm{G} - \overline{\mathrm{G}} \\
\mathbf{P}^t &= \mathrm{T} - \overline{\mathrm{T}}
\end{aligned} \quad (2)
$$

Thus, if all nucleotides have a frequency of $\frac{1}{4}$ (so the vector of means is $[\frac{1}{4}, \frac{1}{4}, \frac{1}{4}, \frac{1}{4}]$, then the simple basis is the set of lines defined by the following vectors:

$$
\begin{matrix} \mathbf{P}^a & \mathbf{P}^c & \mathbf{P}^g & \mathbf{P}^t \\
\begin{bmatrix} \frac{3}{4} \\ -\frac{1}{4} \\ -\frac{1}{4} \\ -\frac{1}{4} \end{bmatrix} &
\begin{bmatrix} -\frac{1}{4} \\ \frac{3}{4} \\ -\frac{1}{4} \\ -\frac{1}{4} \end{bmatrix} &
\begin{bmatrix} -\frac{1}{4} \\ -\frac{1}{4} \\ \frac{3}{4} \\ -\frac{1}{4} \end{bmatrix} &
\begin{bmatrix} -\frac{1}{4} \\ -\frac{1}{4} \\ -\frac{1}{4} \\ \frac{3}{4} \end{bmatrix}
\end{matrix} \quad (3)
$$

#### The Conditional Basis

To conctruct the conditional basis for a site in a sequence, we note that in a system with $n$ mutually exclusive categories, all possible states of the system live on a simplex of dimension $n - 1$. Figure 1A illustrates this for a simplified system of three nucleotides. For the case of 4 nucleotide bases, the simplex is 3 dimensional. This is a tetrahedron with each vertex corresponding to a different nucleotide, as shown in Figure 1B.

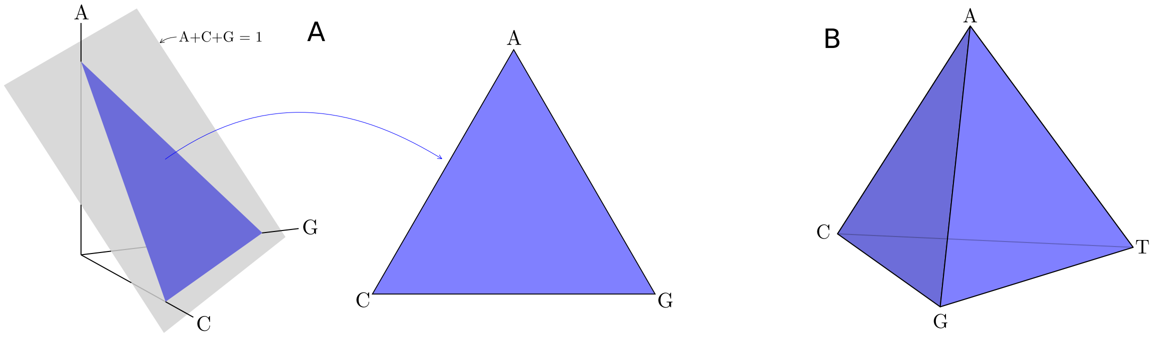


*Figure 1: A) Case of three nucleotides, A, C, and G. Because a particular site on a particular molecule must have one and only one of the possible nucleotides, all possible states of the system lie on the simplex defined by the plane A+C+G=1 and the fact that all values are non-negative. B) With four nucleotides, the simplex is now three dimensional. This simplex is the set of points, in a four dimensional space, in which A+C+G+T = 1 and A,C,G,T ≥ 0.*

We will construct the conditional basis by finding vectors that point towards each of the four vertices of the simplex in Figure 1B and that are orthogonal to the opposite face of the simplex. In other words, instead of finding a vector that is orthogonal to each of the other states, we find one that is orthogonal to the plane that contains them.

Figure 2 shows how we can construct a conditional basis for A, $\mathbf{P}^{a}_{\bullet}$, from the edges of the simplex. We can use any three edges that span the simplex (meaning that at each vertex is is contacted by at least one edge). To construct $\mathbf{P}^{a}_{\bullet}$, we first subtract the mean from each of the three edges, which moves their origins to the population mean (Figure 2B). We next find $\mathbf{P}^{a\cdot c}$ independent of the other two vectors (Figure 2C). The resulting vector, $\mathbf{P}^{a}_{\bullet}$, is orthogonal to the entire face of the simplex that is opposite vertex A. In other words, it captures the effects of A independent of the plane that contains the other nucleotides.

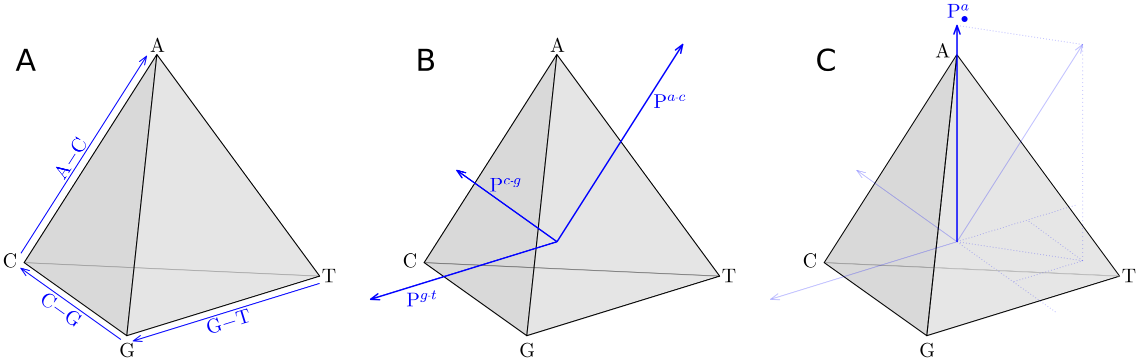


*Figure 2: Constructing a conditional basis. A) We start with three edges that span the simplex; one of which ends at vertex* A*. B) Subtracting the population means from each of these edges yields a basis with the origin at the population mean. C) Calculating* $\mathbf{P}^{a\cdot c}$ *independent of* $\mathbf{P}^{c\cdot g}$ *and* $\mathbf{P}^{g\cdot t}$*, by subtracting it's projection onto them, yields a vector,* $\mathbf{P}^{a}_{\bullet}$*, that captures vertex* A *independent of the plane containing* C, G, *and* T.

Note that, unlike the case for scalar traits, the simple and conditional bases here need not be biorthogonal to one another. To see this, consider the simple case of three possible nucleotides shown in Figure 3. The simple basis ($\mathbf{P}^{a}$, $\mathbf{P}^{c}$, and $\mathbf{P}^{g}$) is shown in Figure 3A along with the simplex within which all actual states of the population reside. Note that the simple basis lies outside of the simplex except for where it touches it at the origin.

The conditional basis ($\mathbf{P}^{a}_{\bullet}$, $\mathbf{P}^{c}_{\bullet}$, and $\mathbf{P}^{g}_{\bullet}$) for this simplified case is shown in Figure 3B. Note that these vectors all lie within the plane of the simplex. The non-biorthogonality of these two different bases can be seen in the fact that $\mathbf{P}^{a}_{\bullet}$ is not orthogonal to $\mathbf{P}^{c}$.

$$\begin{aligned}
\mathbf{P}^{a}_{\bullet} &= \mathbf{P}^{a\cdot c}_{c\cdot g, g\cdot t} \\
\mathbf{P}^{c}_{\bullet} &= \mathbf{P}^{c\cdot g}_{g\cdot t, t\cdot a} \\
\mathbf{P}^{g}_{\bullet} &= \mathbf{P}^{g\cdot t}_{t\cdot a, a\cdot c} \\
\mathbf{P}^{t}_{\bullet} &= \mathbf{P}^{t\cdot a}_{a\cdot c, c\cdot g} \qquad (4)
\end{aligned}$$

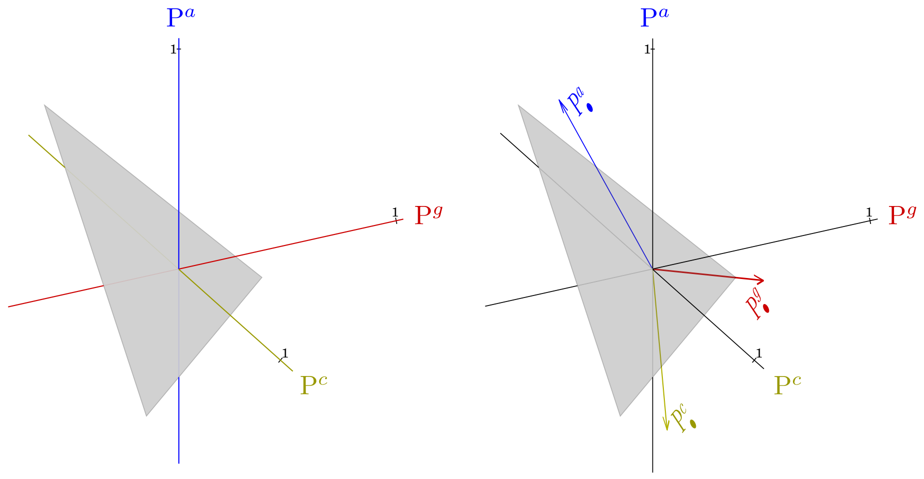


*Figure 3: (A) The simple basis ($\mathbf{P}^a$, $\mathbf{P}^c$, and $\mathbf{P}^g$) for a system with only three states (A, C, and G). The origin is the point on the simplex that corresponds to the frequencies (mean values) of the different states. In this example, A and C each have frequency $\frac{1}{4}$ and G has frequency $\frac{1}{2}$ in the population. (B) The conditional bases, $\mathbf{P}^a_\bullet$, $\mathbf{P}^c_\bullet$, and $\mathbf{P}^g_\bullet$. These bases lie completely within the plane of the simplex.*

Our key result for studying sequence traits is that we can use the simple and conditional bases constructed here just as we used those defined earlier for scalar traits. Thus, for a variable $\phi$, with $\widetilde{\phi}_s$ being the mean value of $\phi$ among individuals with a particular sequence, $s$, we can write:

$$\widetilde{\phi}_s = \overline{\phi}_s + \sum_{i=1}^{n} \left[\!\left[ \begin{matrix} \phi \\ \mathbf{P}^i_\bullet \end{matrix} \right]\!\right] \mathbf{P}^i \tag{5}$$

The fact that we can do this using non-biorthogonal bases is less surprising when we recognize another difference between the scalar and vector cases: In a scalar case with $n$ variables, the space in which they reside was $n$ dimensional, as were the simple and conditional bases. In the vector case that we are considering here, however, $n$ different states live in an $n-1$ dimensional space. Treating this like the scalar case would thus call for the two bases to be $n-1$ dimensional, like the basis defined by $\mathbf{P}^{a\text{-}c}$, $\mathbf{P}^{c\text{-}g}$, and $\mathbf{P}^{g\text{-}t}$ in Figure 2B.

In Equation 5, however, both simple and conditional bases are 4 dimensional, even though all states of the population live in a three dimensional space.

#### Proof

We first construct simple and conditional bases on the simplex, then show that the four conditional polynomials within the simplex form a conditional basis for the four simple polynomials in Equations 2 that lie outside of the simplex.

Since the simplex is three dimensional, we can choose any three edges that span the space to be our simple basis within the simplex. For example, if we use the A-C, C-G, and G-T edges (as in Figure 2A), then subtracting the mean from each yields the simple basis inside the simplex:

$$\begin{aligned} \mathbf{P}^{ac} &= \mathbf{P}^{a} - \mathbf{P}^{c} \\ \mathbf{P}^{cg} &= \mathbf{P}^{c} - \mathbf{P}^{g} \\ \mathbf{P}^{gt} &= \mathbf{P}^{g} - \mathbf{P}^{t} \end{aligned} \tag{6}$$

We seek to show that we can use the conditional basis constructed inside the simplex and the original simple basis which lies outside the simplex (the axes in Figure 3), as we would use the conditional and simple bases in a scalar biorthogonal system (even though these are not biorthogonal).

In other words, we need to show that:

$$\left[\!\!\left[ \begin{matrix} \phi \\ \mathbf{P}^{ac}_{ag,at} \end{matrix} \right]\!\!\right] \mathbf{P}^{ac} + \left[\!\!\left[ \begin{matrix} \phi \\ \mathbf{P}^{ag}_{ac,at} \end{matrix} \right]\!\!\right] \mathbf{P}^{ag} + \left[\!\!\left[ \begin{matrix} \phi \\ \mathbf{P}^{at}_{ac,ag} \end{matrix} \right]\!\!\right] \mathbf{P}^{at} = \left[\!\!\left[ \begin{matrix} \phi \\ \mathbf{P}^{a}_{\bullet} \end{matrix} \right]\!\!\right] \mathbf{P}^{a} + \left[\!\!\left[ \begin{matrix} \phi \\ \mathbf{P}^{c}_{\bullet} \end{matrix} \right]\!\!\right] \mathbf{P}^{c} + \left[\!\!\left[ \begin{matrix} \phi \\ \mathbf{P}^{g}_{\bullet} \end{matrix} \right]\!\!\right] \mathbf{P}^{g} + \left[\!\!\left[ \begin{matrix} \phi \\ \mathbf{P}^{t}_{\bullet} \end{matrix} \right]\!\!\right] \mathbf{P}^{t} \tag{7}$$

The lefthand side of Equation 7 is the basic biorthogonal representation that we used in earlier sections. Three simple bases are chosen that span the simplex, $\mathbf{P}^{ac}$, $\mathbf{P}^{ag}$, and $\mathbf{P}^{at}$, and then conditional bases are found by calculating each simple basis independent of the other two. By the conditional regression theorem (), these are sufficient to describe the linear effects of nucleotide state on another trait. They do not, however, make clear what the independent contribution of each possible nucleotide is. We would thus like to use the righthand side of Equation 7, since this identifies the contribution of each nucleotide.

Preliminaries:

A number of useful relationships follow from the symmetry of the simplex.

1. An edge is equal to any other path between the endpoints of that edge. So, for example:

$$\mathbf{P}^{at} = \mathbf{P}^{ac} + \mathbf{P}^{cg} + \mathbf{P}^{gt} \tag{8}$$

2. Any path from a point back to itself is zero:

$$\mathbf{P}^{ac} + \mathbf{P}^{cg} + \mathbf{P}^{gt} + \mathbf{P}^{ta} = 0 \tag{9}$$

3. Because $\mathbf{P}^{i\cdot j} = -\mathbf{P}^{j\cdot i}$,

$$\mathbf{P}^{ac}_{ag,at} = -\mathbf{P}^{c}_{\bullet} \qquad \mathbf{P}^{ag}_{ac,at} = -\mathbf{P}^{g}_{\bullet} \qquad \mathbf{P}^{at}_{ag,at} = -\mathbf{P}^{t}_{\bullet} \tag{10}$$

4. Because $\mathbf{P}^{i\text{-}j} = \mathbf{P}^{i} - \mathbf{P}^{j}$, it follows that $\mathbf{P}^{i\text{-}j} + \mathbf{P}^{j\text{-}k} = \mathbf{P}^{i\text{-}k}$. Combined with the fact that $\llbracket a+b \rrbracket = \llbracket a \rrbracket + \llbracket b \rrbracket + 2\llbracket a, b \rrbracket$, we can write some covariances between bases in terms of variances:

$$\left\llbracket \mathbf{P}^{i\text{-}j}, \mathbf{P}^{j\text{-}k} \right\rrbracket = \frac{1}{2}\left( \left\llbracket \mathbf{P}^{i\text{-}k} \right\rrbracket - \left\llbracket \mathbf{P}^{i\text{-}j} \right\rrbracket - \left\llbracket \mathbf{P}^{j\text{-}k} \right\rrbracket \right) \tag{11}$$

Using the fact that $\mathbf{P}^{ij} = \mathbf{P}^{i} - \mathbf{P}^{j}$, and the identities in 10, the lefthand side of Equation 7 can be written:

$$\begin{aligned}
&\left\llbracket \begin{matrix}\phi \\ \mathbf{P}^{ac}_{ag,at}\end{matrix} \right\rrbracket \mathbf{P}^{ac} + \left\llbracket \begin{matrix}\phi \\ \mathbf{P}^{ag}_{ac,at}\end{matrix} \right\rrbracket \mathbf{P}^{ag} + \left\llbracket \begin{matrix}\phi \\ \mathbf{P}^{at}_{ac,ag}\end{matrix} \right\rrbracket \mathbf{P}^{at} \\
&= \left\llbracket \begin{matrix}\phi \\ \mathbf{P}^{ac}_{ag,at}\end{matrix} \right\rrbracket \mathbf{P}^{a} + \left\llbracket \begin{matrix}\phi \\ \mathbf{P}^{ag}_{ac,at}\end{matrix} \right\rrbracket \mathbf{P}^{a} + \left\llbracket \begin{matrix}\phi \\ \mathbf{P}^{at}_{ac,ag}\end{matrix} \right\rrbracket \mathbf{P}^{a} + \left\llbracket \begin{matrix}\phi \\ \mathbf{P}^{c}_{\bullet}\end{matrix} \right\rrbracket \mathbf{P}^{c} + \left\llbracket \begin{matrix}\phi \\ \mathbf{P}^{g}_{\bullet}\end{matrix} \right\rrbracket \mathbf{P}^{g} + \left\llbracket \begin{matrix}\phi \\ \mathbf{P}^{t}_{\bullet}\end{matrix} \right\rrbracket \mathbf{P}^{t}
\end{aligned} \tag{12}$$

In order to confirm Equation 7, we thus need to show that:

$$\left\llbracket \begin{matrix}\phi \\ \mathbf{P}^{ac}_{ag,at}\end{matrix} \right\rrbracket + \left\llbracket \begin{matrix}\phi \\ \mathbf{P}^{ag}_{ac,at}\end{matrix} \right\rrbracket + \left\llbracket \begin{matrix}\phi \\ \mathbf{P}^{at}_{ac,ag}\end{matrix} \right\rrbracket = \left\llbracket \begin{matrix}\phi \\ \mathbf{P}^{a}_{\bullet}\end{matrix} \right\rrbracket \tag{13}$$

Using the identities in 10, this is the same as saying that:

$$\left\llbracket \begin{matrix}\phi \\ \mathbf{P}^{a}_{\bullet}\end{matrix} \right\rrbracket + \left\llbracket \begin{matrix}\phi \\ \mathbf{P}^{c}_{\bullet}\end{matrix} \right\rrbracket + \left\llbracket \begin{matrix}\phi \\ \mathbf{P}^{g}_{\bullet}\end{matrix} \right\rrbracket + \left\llbracket \begin{matrix}\phi \\ \mathbf{P}^{t}_{\bullet}\end{matrix} \right\rrbracket = 0 \tag{14}$$

Since the regression of anything with a constant is zero, Equation 14 must hold if:

$$\frac{\mathbf{P}^{a}_{\bullet}}{\llbracket \mathbf{P}^{a}_{\bullet} \rrbracket} + \frac{\mathbf{P}^{c}_{\bullet}}{\llbracket \mathbf{P}^{c}_{\bullet} \rrbracket} + \frac{\mathbf{P}^{g}_{\bullet}}{\llbracket \mathbf{P}^{g}_{\bullet} \rrbracket} + \frac{\mathbf{P}^{t}_{\bullet}}{\llbracket \mathbf{P}^{t}_{\bullet} \rrbracket} = \text{constant} \tag{15}$$

Confirming Equation 15 will confirm Equation 7 and our result from 5. In order to confirm (15), we need to deal with the variances in the denominators. A way to do this is suggested by a symmetry of the covariance matrix.

In Equations 4, we used the edges $(A - C)$, $(C - G)$, $(G - T)$, and $(T - A)$ to define the conditional bases. The covariance matrix for the four corresponding bases is:

$$\mathbb{C} = \begin{bmatrix}
\llbracket \mathbf{P}^{a\text{-}c} \rrbracket & \llbracket \mathbf{P}^{a\text{-}c}, \mathbf{P}^{c\text{-}g} \rrbracket & \llbracket \mathbf{P}^{a\text{-}c}, \mathbf{P}^{g\text{-}t} \rrbracket & \llbracket \mathbf{P}^{a\text{-}c}, \mathbf{P}^{t\text{-}a} \rrbracket \\
\llbracket \mathbf{P}^{c\text{-}g}, \mathbf{P}^{a\text{-}c} \rrbracket & \llbracket \mathbf{P}^{c\text{-}g} \rrbracket & \llbracket \mathbf{P}^{c\text{-}g}, \mathbf{P}^{g\text{-}t} \rrbracket & \llbracket \mathbf{P}^{c\text{-}g}, \mathbf{P}^{t\text{-}a} \rrbracket \\
\llbracket \mathbf{P}^{g\text{-}t}, \mathbf{P}^{a\text{-}c} \rrbracket & \llbracket \mathbf{P}^{g\text{-}t}, \mathbf{P}^{c\text{-}g} \rrbracket & \llbracket \mathbf{P}^{g\text{-}t} \rrbracket & \llbracket \mathbf{P}^{g\text{-}t}, \mathbf{P}^{t\text{-}a} \rrbracket \\
\llbracket \mathbf{P}^{t\text{-}a}, \mathbf{P}^{a\text{-}c} \rrbracket & \llbracket \mathbf{P}^{t\text{-}a}, \mathbf{P}^{c\text{-}g} \rrbracket & \llbracket \mathbf{P}^{t\text{-}a}, \mathbf{P}^{g\text{-}t} \rrbracket & \llbracket \mathbf{P}^{t\text{-}a} \rrbracket
\end{bmatrix} \tag{16}$$

For each basis, we used only three edges (because the system is three dimensional). Thus, the calculation of any one basis vector used only terms corresponding to a submatrix of (16) obtained by removing the row and column corresponding to one of the bases. For example, the calculation of $\mathbf{P}^{a}_{\bullet}$ in Equations 4 did not include $\mathbf{P}^{t\text{-}a}$.

This matters because the matrix in 16 is the covariance matrix for a multinomial distribution (in which a given site can be in one and only one of four mutually exclusive states). The determinant of the entire matrix $\mathbb{C}$ is zero. However, May & Johnson (1998) show that the determinants of the four submatrices resulting from eliminating the row and column corresponding to each basis are not zero, and furthermore are the same.

Using the corollary to the variance-determinant theorem (Equation 26 in the other document), we can write the determinant of these submatrices in terms of the variances of the orthogonal polynomials derived from the edges of the simplex. Since the determinants of all of these submatrices are the same, we have:

$$\begin{aligned} |\mathbb{C}_{sub}| &= [\![\mathbf{P}^{a}_{\bullet}]\!] \left[\!\!\left[\mathbf{P}^{c\text{-}g}_{g\text{-}t}\right]\!\!\right] [\![\mathbf{P}^{g\text{-}t}]\!] \\ &= [\![\mathbf{P}^{c}_{\bullet}]\!] \left[\!\!\left[\mathbf{P}^{g\text{-}t}_{t\text{-}a}\right]\!\!\right] [\![\mathbf{P}^{t\text{-}a}]\!] \\ &= [\![\mathbf{P}^{g}_{\bullet}]\!] [\![\mathbf{P}^{t\text{-}a}_{a\text{-}c}]\!] [\![\mathbf{P}^{a\text{-}c}]\!] \\ &= [\![\mathbf{P}^{t}_{\bullet}]\!] \left[\!\!\left[\mathbf{P}^{a\text{-}c}_{c\text{-}g}\right]\!\!\right] [\![\mathbf{P}^{c\text{-}g}]\!] \end{aligned} \tag{17}$$

Multiplying both sides of Equation 15 by $|\mathbb{C}_{sub}|$ gives us the condition:

$$\mathbf{P}^{a}_{\bullet} \left[\!\!\left[\mathbf{P}^{c\text{-}g}_{g\text{-}t}\right]\!\!\right] [\![\mathbf{P}^{g\text{-}t}]\!] + \mathbf{P}^{c}_{\bullet} \left[\!\!\left[\mathbf{P}^{g\text{-}t}_{t\text{-}a}\right]\!\!\right] [\![\mathbf{P}^{t\text{-}a}]\!] + \mathbf{P}^{g}_{\bullet} [\![\mathbf{P}^{t\text{-}a}_{a\text{-}c}]\!] [\![\mathbf{P}^{a\text{-}c}]\!] + \mathbf{P}^{t}_{\bullet} \left[\!\!\left[\mathbf{P}^{a\text{-}c}_{c\text{-}g}\right]\!\!\right] [\![\mathbf{P}^{c\text{-}g}]\!] = \text{constant} \tag{18}$$

To confirm Equation 15, we will show that the lefthand side can be written as:

$$\left(\mathbf{P}^{a\text{-}c} + \mathbf{P}^{c\text{-}g} + \mathbf{P}^{g\text{-}t} + \mathbf{P}^{t\text{-}a}\right) \cdot \gamma \tag{19}$$

where $\gamma$ is determined by the frequencies of the different nucleotides. Combined with Equation 9, this will confirm Equation 15 and show that the constant is in fact 0.

We start by identifying all terms that multiply $\mathbf{P}_{\bullet}ac$ in Equation 18. $\mathbf{P}_{\bullet}ac$ appears in the calculation of $\mathbf{P}^{a}_{\bullet}$, $\mathbf{P}^{g}_{\bullet}$, and $\mathbf{P}^{t}_{\bullet}$. It does not appear in the calculation of $\mathbf{P}^{c}_{\bullet}$. With this in mind, we find the parts of Equation 18 that include $\mathbf{P}^{a\text{-}c}$ to be:

$$\mathbf{P}^{a\text{-}c} \left( \left[\!\!\left[\mathbf{P}^{c\text{-}g}_{g\text{-}t}\right]\!\!\right] [\![\mathbf{P}^{g\text{-}t}]\!] + \left[\!\!\left[\begin{matrix}\mathbf{P}^{g\text{-}t} \\ \mathbf{P}^{t\text{-}a}_{a\text{-}c}\end{matrix}\right]\!\!\right] \left[\!\!\left[\begin{matrix}\mathbf{P}^{t\text{-}a} \\ \mathbf{P}^{a\text{-}c}\end{matrix}\right]\!\!\right] [\![\mathbf{P}^{t\text{-}a}_{a\text{-}c}]\!] [\![\mathbf{P}^{a\text{-}c}]\!] - \left[\!\!\left[\begin{matrix}\mathbf{P}^{g\text{-}t} \\ \mathbf{P}^{a\text{-}c}\end{matrix}\right]\!\!\right] [\![\mathbf{P}^{t\text{-}a}_{a\text{-}c}]\!] [\![\mathbf{P}^{a\text{-}c}]\!] - \left[\!\!\left[\begin{matrix}\mathbf{P}^{t\text{-}a} \\ \mathbf{P}^{a\text{-}c}_{c\text{-}g}\end{matrix}\right]\!\!\right] \left[\!\!\left[\mathbf{P}^{a\text{-}c}_{c\text{-}g}\right]\!\!\right] [\![\mathbf{P}^{c\text{-}g}]\!] \right) \tag{20}$$

If the terms in the parentheses in Equation 20 are equal to some symmetrical value of the simplex – meaning that they are not just a consequence of starting with the $(A - C)$ edge – then we can conclude that we would get the same value for any edge. This would confirm Equation 19 and prove our result.

Using the rule in Equation 11, we can write the terms in parentheses in expression 20 completely in terms of variances of edges of the simplex. Doing this, we find that this expression is equal to:

$$\frac{1}{4}\Big(\llbracket \mathbf{P}^{c\cdot g}\rrbracket \llbracket \mathbf{P}^{g\cdot t}\rrbracket + \llbracket \mathbf{P}^{c\cdot t}\rrbracket \llbracket \mathbf{P}^{t\cdot g}\rrbracket + \llbracket \mathbf{P}^{g\cdot a}\rrbracket \llbracket \mathbf{P}^{a\cdot t}\rrbracket + \llbracket \mathbf{P}^{a\cdot t}\rrbracket \llbracket \mathbf{P}^{t\cdot c}\rrbracket + \llbracket \mathbf{P}^{t\cdot a}\rrbracket \llbracket \mathbf{P}^{a\cdot c}\rrbracket + \llbracket \mathbf{P}^{g\cdot t}\rrbracket \llbracket \mathbf{P}^{t\cdot a}\rrbracket$$
$$+ \llbracket \mathbf{P}^{a\cdot c}\rrbracket \llbracket \mathbf{P}^{c\cdot g}\rrbracket + \llbracket \mathbf{P}^{a\cdot g}\rrbracket \llbracket \mathbf{P}^{g\cdot t}\rrbracket + \llbracket \mathbf{P}^{a\cdot c}\rrbracket \llbracket \mathbf{P}^{c\cdot t}\rrbracket + \llbracket \mathbf{P}^{c\cdot a}\rrbracket \llbracket \mathbf{P}^{a\cdot g}\rrbracket + \llbracket \mathbf{P}^{g\cdot c}\rrbracket \llbracket \mathbf{P}^{c\cdot t}\rrbracket + \llbracket \mathbf{P}^{c\cdot g}\rrbracket \llbracket \mathbf{P}^{g\cdot a}\rrbracket$$
$$- \llbracket \mathbf{P}^{a\cdot c}\rrbracket^2 - \llbracket \mathbf{P}^{a\cdot g}\rrbracket^2 - \llbracket \mathbf{P}^{a\cdot t}\rrbracket^2 - \llbracket \mathbf{P}^{c\cdot g}\rrbracket^2 - \llbracket \mathbf{P}^{c\cdot t}\rrbracket^2 - \llbracket \mathbf{P}^{g\cdot t}\rrbracket^2\Big) \quad (21)$$

The terms in (21) are just the sum of products of the variances of all adjacent edges of the simplex, minus the sum of squared variances of each edge. Since this value involves all edges equally, it should be the same regardless of which one we start with. We thus conclude that Equation 19 is true, with $\gamma$ being equal to the expression in (21. It follows that Equation 7 is true.
